# The C-terminal helix of BubR1 is essential for CENP-E-dependent chromosome alignment

**DOI:** 10.1101/2020.02.25.962613

**Authors:** Thibault Legal, Daniel Hayward, Agata Gluszek-Kustusz, Elizabeth A. Blackburn, Christos Spanos, Juri Rappsilber, Ulrike Gruneberg, Julie P.I. Welburn

**Affiliations:** Wellcome Trust Centre for Cell Biology, School of Biological Sciences, University of Edinburgh, Edinburgh EH9 3BF, Scotland, UK; Sir William Dunn School of Pathology, University of Oxford, South Parks Road, Oxford, UK; Chair of Bioanalytics, Institute of Biotechnology, Technische Universität Berlin, Berlin, German

**Keywords:** CENP-E, motor, kinetochore, mitosis, microtubule

## Abstract

During cell division, misaligned chromosomes are captured and aligned by motors before their segregation. The CENP-E motor is recruited to polar unattached kinetochores, to facilitate chromosome alignment. The spindle checkpoint protein BubR1 has been reported as a CENP-E interacting partner, but to what extent, if at all, BubR1 contributes to CENP-E localization at kinetochores, has remained controversial. Here we define the molecular determinants that specify the interaction between BubR1 and CENP-E. The basic C-terminal helix of BubR1 is necessary but not sufficient for CENP-E interaction, while a minimal key acidic patch on the kinetochore-targeting domain of CENP-E, is also essential. We then demonstrate that BubR1 is required for the recruitment of CENP-E to kinetochores to facilitate chromosome alignment. This BubR1-CENP-E axis is critical to align chromosomes that have failed to congress through other pathways and recapitulates the major known function of CENP-E. Overall, our studies define the molecular basis and the function for CENP-E recruitment to BubR1 at kinetochores during mammalian mitosis.

## Introduction

To maintain their genomic integrity, eukaryotic cells must distribute their DNA equally to the daughter cells. Spindle microtubules mediate the segregation of chromosomes, by associating with the kinetochore of chromosomes, a large protein complex that mediates the end-on attachment of chromosomes to microtubules. At mitotic onset, chromosomes are dispersed throughout the cytoplasm, posing a challenge for their capture by microtubules from opposite poles, a pre-requisite for their accurate segregation. Multiple pathways involving microtubules and motors co-exist to ensure chromosome congression and bi-orientation (Maiato et al., 2017). A subset of chromosomes that lie outside the interpolar region assembling the spindle are dependent on CENP-E for congression. CENP-E is a large 312 kDa plus end-directed kinesin that is recruited to unattached and unaligned kinetochores, and to the outer corona, that expands around kinetochores to maximize microtubule capture (Cooke et al., 1997; Yao et al., 1997). Kinetochore-bound CENP-E moves laterally attached chromosomes to the cell equator along microtubules (Wood et al., 1997). CENP-E may also help sort kinetochore-nucleated microtubules and promote end-on attachments and biorientation (Shrestha and Draviam, 2013; Sikirzhytski et al., 2018). CENP-E then remains at aligned kinetochores, albeit with lower levels, where it plays a role in maintaining a robust connection between kinetochores and microtubules during metaphase, and during anaphase as the kinetochores are pulled to opposite poles by depolymerizing microtubules (Brown et al., 1996; Vitre et al., 2014).

CENP-E is enriched at unattached and misaligned kinetochores in early mitosis. The human CENP-E kinetochore-targeting domain has previously been mapped (Chan et al., 1998). Over the years, CENP-E has been reported to interact with multiple kinetochore proteins: BubR1, CENP-F, Clasp2, Mad1, and other interactors such as Septin, CKAP5, NPM1 (Akera et al., 2015; Chan et al., 1998; Maffini et al., 2009; Zhu et al., 2008; Maliga et al., 2013). Post-translational modifications may also enhance CENP-E targeting to kinetochores (Ashar et al., 2000; Zhang et al., 2008). Overall, the molecular basis for CENP-E recruitment to kinetochores remains poorly understood. Its recruitment there through a dependency with the spindle-checkpoint proteins, budding uninhibited by benzimidazole 1 (Bub1) and Bub1-related (BubR1) mitotic checkpoint Ser/Thr kinases has been shown; in *Xenopus* and DLD-1 cells, CENP-E kinetochore levels are strongly reduced upon BubR1 depletion (Johnson et al., 2004; Mao et al., 2003). Other studies however argue CENP-E levels are not affected by BubR1 depletion. (Chan et al., 1999; Ciossani et al., 2018; Kops et al., 2004). These observed differences could be due to distinct experimental setups and different cell types and species. CENP-E recruitment to the outer corona appears independent of CDK1 activity but depends on the presence of the RZZ complex (Pereira et al., 2018; Sacristan et al., 2018). In the absence of the RZZ complex, CENP-E is recruited to kinetochores but not to the expandable corona. CENP-E remains at bioriented kinetochores after removal of checkpoint proteins, disassembly of the outer corona and throughout anaphase, indicating CENP-E has multiple yet unidentified binding partners at the kinetochore (Brown et al., 1996; Cooke et al., 1997; Gudimchuk et al., 2013). Here we characterized the kinetochore targeting domain of CENP-E biophysically and used a non-biased approach to find mitotic partners of CENP-E. We found BubR1 as a major interactor for this domain of CENP-E and defined the molecular requirements for the BubR1-CENP-E interaction. Overall, we demonstrate BubR1 contributes to CENP-E localization to kinetochores at aligned chromosomes and during spindle checkpoint activation and that this axis is essential for facilitating chromosome alignment.

## Results

To define the regulation of CENP-E targeting to kinetochores, we examined quantitatively endogenous CENP-E levels at kinetochores during distinct stages of cell division, CENP-E levels were maximal during prometaphase and decreased during metaphase (Fig S1A, B). We also observed endogenous CENP-E on microtubules in early stages of mitosis, although microtubules did not appear necessary for its kinetochore targeting. Indeed CENP-E levels were largely comparable to that in prometaphase upon nocodazole-induced depolymerization of microtubules, creating unattached kinetochores (Fig S1A, B). To analyze the molecular requirements for CENP-E localization, we precisely mapped the regions of CENP-E isoform 1 that target to kinetochores and centrosomes using transient transfection of CENP-E constructs fused to GFP (Fig 1A). CENP-E_2055-2608_ in the C terminus of CENP-E, largely similar to the previously published kinetochore-targeting construct 1958-2628, was necessary and sufficient for targeting to kinetochores in HeLa cells (Fig 1B, C)(Chan et al., 1998). The shorter CENP-E_2055-2450_ still showed kinetochore localization, while an even shorter CENP-E_2055-2356_ targeted weakly to a subset of kinetochores (Fig 1B, C). This heterogeneous targeting was previously observed and is likely to reflect different attachment states or kinetochore heterogeneity (Chan et al., 1998). In the absence of the first 35 amino acids in this domain, CENP-E_2090-2450_ lost the ability to target to kinetochores (Fig 1B, C). However, CENP-E_2260-2608_ localized specifically to a region between the two centrioles or associated closely with one centriole both in interphase and mitosis (Fig S1C). The intercentriole interacting proteins remain unknown and this interaction will not be pursued further here.

**Figure 1:**
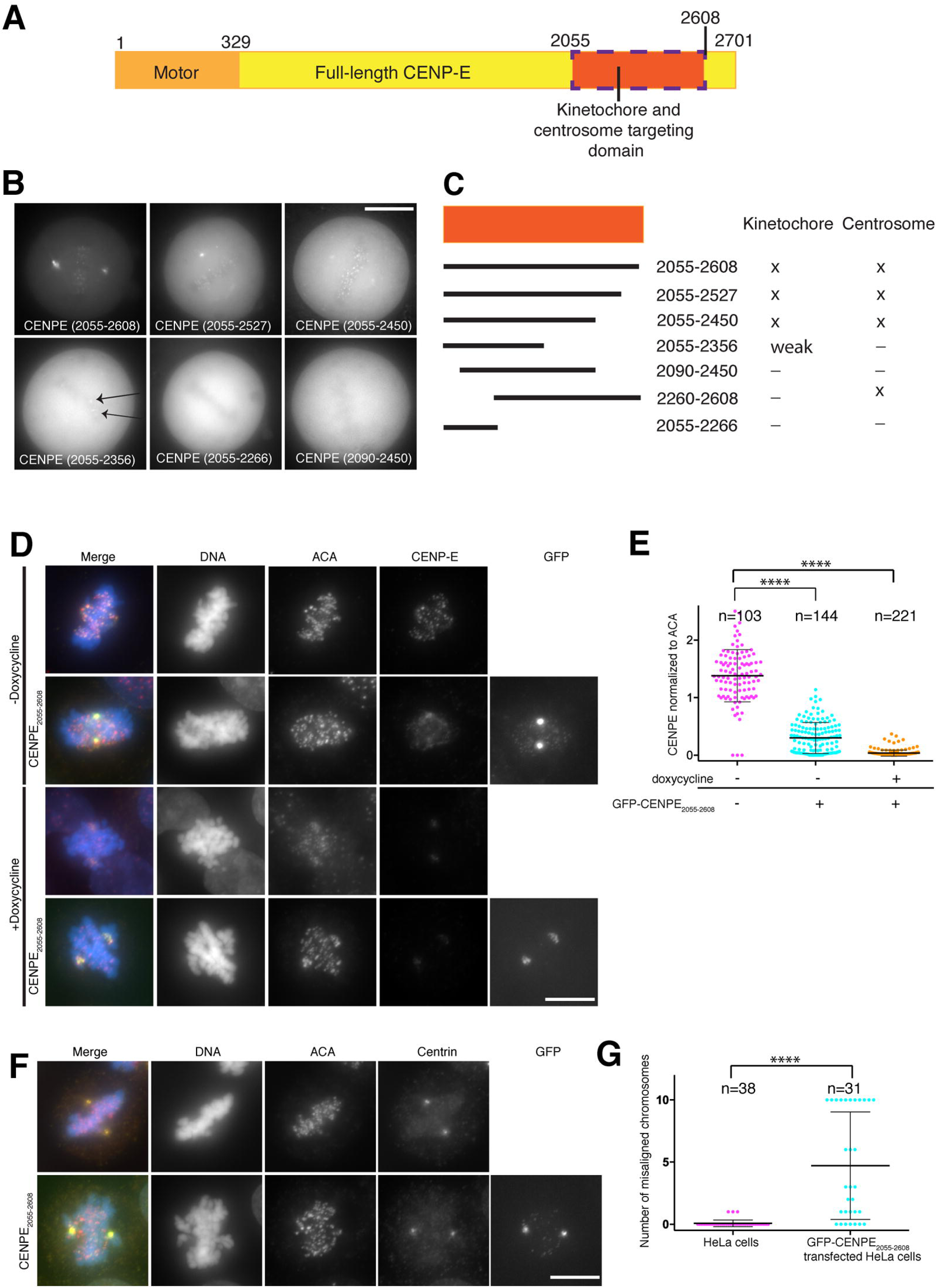
Mapping of the kinetochore- and centrosome-targeting domain of CENP-E. (A) Schematic diagram of CENP-E, highlighting the motor (apricot) and kinetochore- and centrosome-targeting domains (orange). (B) Representative images of live HeLa cells transfected with GFP-CENP-E constructs (scale bar: 10 μm) and (C) representative map of the corresponding kinetochore and centrosome targeting domains for human CENP-E. (D) Representative immunofluorescence images of HeLa cells transfected with GFP-CENP-E_2055-2608_ in the presence and knockout of endogenous CENP-E and stained for endogenous CENP-E, ACA and DNA. Scale bar: 10 μm. (E) Scatter dot plot showing quantification of CENP-E intensity normalized to ACA, in the presence and knockout of endogenous CENP-E and GFP-CENP-E_2055-2608_. Each point represents the intensity of CENP-E over ACA at one kinetochore. Black line represents the mean and whiskers represent the standard deviation. Asterisks indicate ordinary One-way Anova test significance value. ****P<0.0001. (F) Representative immunofluorescence images of HeLa cells transfected with GFP-CENP-E_2055-2608_ and stained for centrin, ACA and for DNA. Scale bar: 10 μm. (G) Scatter dot plot showing the number of misaligned chromosomes in HeLa cells and GFP-CENP-E_2055-2608_ transfected HeLa cells. Each point represents one cell with corresponding number of misaligned chromosomes. Black line represents the mean and whiskers represent the standard deviation. Asterisks indicate Unpaired T-test significance value. ****P<0.0001.

To further define how CENP-E targets to kinetochores, we tested whether CENP-E_2055-2608_ dimerizes with endogenous CENP-E at kinetochores. We depleted CENP-E using a Cas9 inducible cell line expressing a CENP-E sgRNA (McKinley and Cheeseman, 2017). CENP-E was largely depleted after 72 hours (Fig 1D, E; + doxycyline). All CENP-E depleted cells displayed issues with chromosome alignment although the levels of CENP-E depletion varied from cell to cell. Cells clearly depleted for endogenous CENP-E, identified by immunofluorescence, were analyzed. In the absence of endogenous CENP-E, GFP-CENP-E_2055-2608_ was only weakly targeted to most kinetochores, indicating that CENP-E_2055-2608_ recruitment to kinetochores may depend on the full-length endogenous CENP-E or that CENP-E removal affects other kinetochore proteins necessary for its recruitment (Fig 1D, S1D). However, GFP-CENP-E_2055-2608_ was strongly enriched at kinetochores close to spindle poles, suggesting GFP-CENP-E_2055-2608_ was recruited to these kinetochore subpopulations independently of endogenous CENP-E, through another binding partner (Fig 1D, S1E). GFP-CENP-E_2055-_ 2608 appeared to compete with CENP-E at kinetochores, causing a reduction in endogenous CENP-E at kinetochores (Fig 1E) in agreement with (Schaar et al., 1997). Additionally, CENP-E_2055-2608_ transfection caused many chromosomes to become misaligned (Fig 1D, F, G), presumably by replacing endogenous motor-domain containing CENP-E at kinetochores. Overall these results indicate CENP-E has distinct spindle pole- and kinetochore-targeting domains close to each other. CENP-E_2055-2608_ targeting to kinetochores outcompetes endogenous CENP-E but it may also favor the targeting of kinetochores to spindle poles through its spindle pole-targeting region.

To define the molecular basis for the CENP-E kinetochore-targeting domain, we expressed and purified recombinant CENP-E_2055-2608_, which robustly targets to both kinetochores and centrosomes, and CENP-E_2055-2358_, which targets weakly to kinetochores. CENP-E_2055-2608_ aggregated in 150mM NaCl and was maintained in 500mM NaCl. SEC-MALS analysis revealed that CENP-E_2055-2608_ assembles as a dimer in solution, while the minimal kinetochore-targeting domain CENP-E_2055-2358_ was monomeric (Fig S2A, B). Circular dichroism further defined the secondary structural elements of CENP-E_2055-2608_.and CENP-E_2055-2358_ (Fig S2C). CENP-E_2055-2608_ has a α-helical content of around 50.9%, while the shorter domain CENP-E_2055-2358_ is 80% α-helical, with 8.8% containing turns and 12.1% containing unstructured regions (Fig S2D). Thus the region 2358-2608 responsible for dimerization and centrosome region-targeting, is highly likely an α-helical coiled coil. Rotary shadowing further revealed that CENP-E_2055-2608_ is an elongated domain with a globular region at one end and rod-like shape, supporting a coiled coil conformation (Fig S2E). Overall, these data indicated that the CENP-E_2055-2608_ domain can be subdivided into an N-terminal monomeric α-helical rich domain, essential for kinetochore targeting and a C-terminal domain that provides dimerization properties.

We then sought to define the major CENP-E interactors at kinetochores. CENP-E is strongly recruited to unattached kinetochores at mitotic onset (Yen et al., 1991). To identify mitotic interactors of the CENP-E kinetochore-targeting domain, we incubated CENP-E_2055-2358_ with clarified mitotic cell lysate, from cells arrested with the microtubule-depolymerizing drug nocodazole. We then pulled down CENP-E_2055-2358_ and associated proteins, which were subjected to mass spectrometry for identification (Fig S3A, table S1). We found CENP-E_2055-2358_ specifically interacted with BubR1 and MYPT1, a protein phosphatase 1 regulatory subunit 12A (Fig S3B). We did not pursue the interaction of MYPT1 in this study. To test whether CENP-E binds directly BubR1, we expressed and purified recombinant BubR1 from insect cells and analyzed whether stoichiometric amounts of BubR1 could interact with the longer dimeric kinetochore-targeting domain CENP-E_2055-2608_ by size-exclusion chromatography. Indeed full-length BubR1 interacts with CENP-E_2055-2608_ *in vitro* (data not shown). To test whether CENP-E_2055-2608_ interacts with the N terminus of BubR1 or with the pseudokinase domain, we generated two BubR1 constructs containing either the N- or C-terminal domains. The N terminus of BubR1_1-484_ did not co-migrate with CENP-E_2055-2608_ (Fig S3C) while the C terminus containing the pseudokinase domain BubR1_432-1050_ did (Fig S3D). Thus these data indicate CENP-E_2055-2608_ binds to the pseudokinase domain of BubR1. While we were conducting these experiments, a parallel study also reported an interaction between CENP-E and the pseudokinase domain of BubR1 (Ciossani et al., 2018).

Our *in vivo* work indicated CENP-E_2055-2356_ fused to GFP could still weakly associate with kinetochores (Fig 1B). To test whether post-translational modifications were necessary for the interaction, we tested if our recombinant construct CENP-E_2055-_ 2358 could interact with bacterially expressed BubR1_705-1050_ *in vitro*. Indeed CENP-E_2055-2358_ co-eluted with BubR1_705-1050_ indicating they interact in the absence of post-translational modifications (Fig S4A). CENP-E_2055-2358_ is monomeric (Fig S2A, B) and not very stable in low salt. To stabilize it while mimicking dimeric CENP-E_2055-2608_, we fused it to a C-terminal GST and removed 14 residues at the N terminus, to stabilize it while mimicking dimeric CENP-E_2055-2608._ CENP-E_2069-2358_-GST was more stable in low salt concentration and could then be further used to analyze the BubR1-CENP-E interaction by SEC. CENP-E_2069-2358_–GST co-eluted with BubR1_705-1050_, as shown by the shift in the elution profile (Fig 2A), while GST alone did not (Fig S4B). The constructs were monodisperse but we were not able to obtain diffracting crystals of the kinetochore-targeting domain of CENP-E_2055-2358_ alone or bound to BubR1.

**Figure 2:**
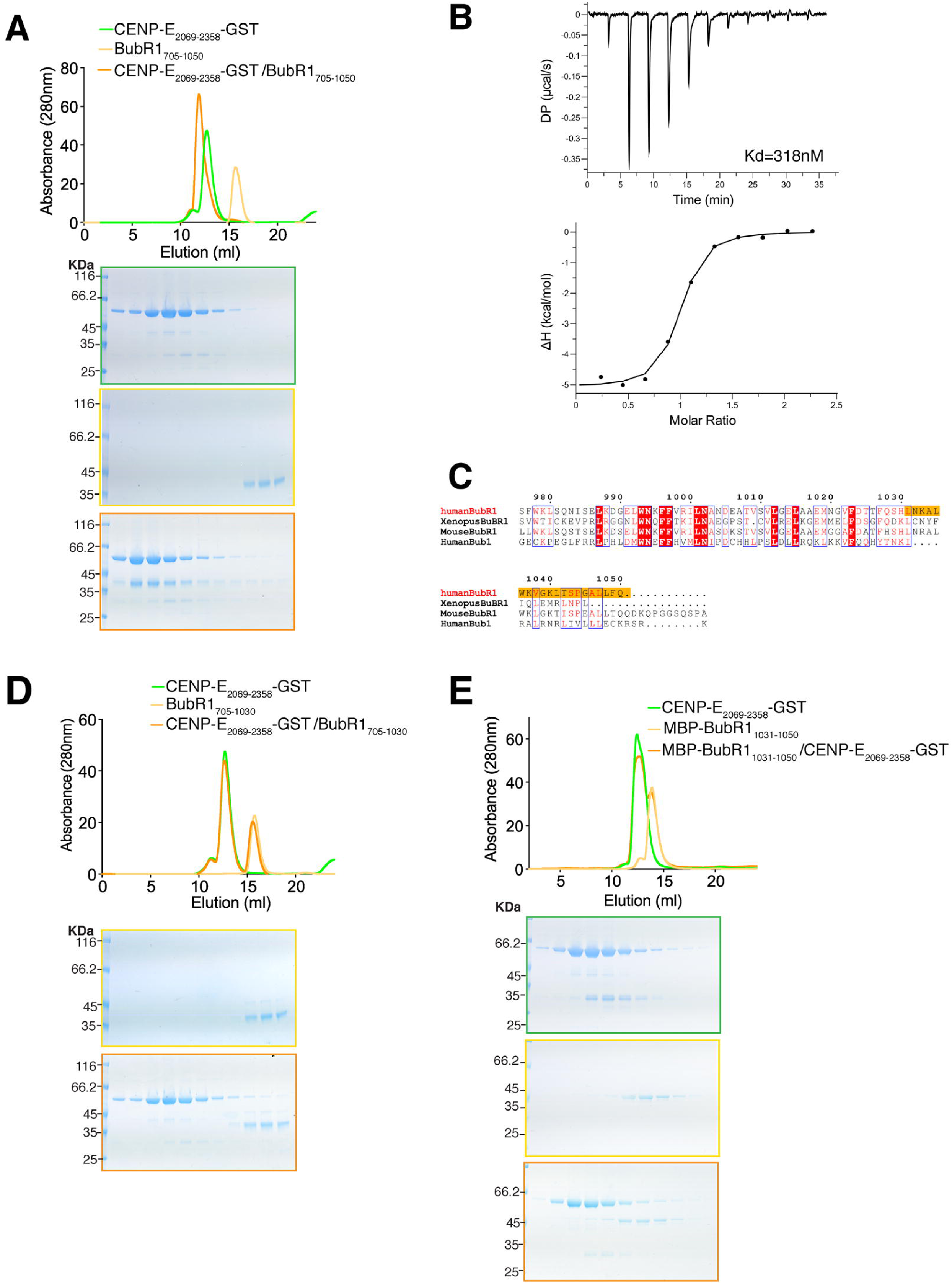
Requirement of the C-terminal helix of BubR1 for CENP-E binding. (A) Top, SEC analysis and elution profile for CENP-E_2069-2358_-GST (green), BubR1_705-1050_ (yellow) and CENP-E_2069-2358_-GST/BubR1_705-1050_ (orange). Bottom, Coomassie-stained gels showing elution profiles for the corresponding protein complexes. (B) Thermodynamics of BubR1_705-1050_/CENP-E_2091-2358_-GST interaction determined by isothermal titration calorimetry. The y-axis indicates kcal/mol of injectant. The dissociation constant (K_d_) between BubR1_705-1050_ and CENP-E_2091-2358_-GST was determined to be 318 ± 90 nM. (C) Sequence alignment of the C terminus of human BubR1 with mouse and *Xenopus* BubR1, and human Bub1. Boxed red and blue are the conserved and similar amino acids across all 4 proteins, respectively. Amino acids in red are those with conserved properties in at least 3 sequences. The sequence necessary for BubR1 binding to CENP-E_2055-2608_ is highlighted in orange. (D, E) SEC analysis and elution profile for CENP-E_2069-2358_-GST (green), BubR1_705-1050_ and MBP-BubR1_1031-1050_ (yellow, C and D respectively), and CENP-E_2069-2358_-GST/BubR1 constructs (orange). Bottom, Coomassie-stained gels showing elution profiles for the corresponding protein complexes.

To investigate the thermodynamics of the BubR1/CENP-E interaction, we performed isothermal titration calorimetry (ITC). The CENP-E_2069-2358_-GST construct had to be optimized slightly to remove some GST contaminants and degradation. We therefore removed a further 22 residues at the N terminus and purified it in complex with BubR1 before separating the complex in high ionic strength using gel filtration. This way, we obtained >95% pure BubR1_705-1050_ and CENP-E_2091-2358_-GST. At 37°C the pseudokinase domain of BubR1 bound CENP-E_2091-2358_-GST with mid-nanomolar affinity with a K_d_=318 ± 90 nM, (Fig 2B). At this temperature, formation of the complex had an exothermic heat signature. The enthalpic and entropic components driving the interaction were of a similar magnitude (ΔH= −5.1 ± 0.2 kcal/mol; −TΔS= −4.1 kcal/mol). The stoichiometry between BubR1 and CENP-E_2091-2358_ was determined to be 1:1. The stoichiometry must be put in the context of full-length dimeric CENP-E. Thus the CENP-E motor is able to bind two molecules of BubR1.

To further map the interaction region specific to BubR1 and CENP-E, we examined the sequence conservation between Bub1 and BubR1 kinase domains. We found a longer loop in the C terminus of BubR1 when compared to Bub1 that showed sequence divergence between human Bub1 and BubR1, but displayed sequence similarity across BubR1 species (Fig 2C). We hypothesized this region may be important for the CENP-E-BubR1 interaction Indeed, CENP-E_2069-2358_-GST did not co-elute with BubR1_705-1030_ lacking the last 20 amino acids (Fig 2D) suggesting that this part of BubR1 is critical for the interaction with CENP-E.. However, on its own this basic helix in BubR1_1030-1050_ (pI=10.30), was not sufficient to interact with CENP-E (Fig 2E). Based on the basic properties of this helix, we also mapped the interaction of BubR1 with CENP-E to the C terminus of CENP-E_2055-2358_. We found a negatively charged region in CENP-E, which we hypothesized could interact with the basic helix of the kinase domain of BubR1. We mutated 4 highly conserved glutamates (E2313, E2316, E2318 and E2319) to alanines in CENP-E_2069-2358_-GST, named thereafter CENP-E_4E_-GST (Fig 3A). CENP-E_4E_-GST was co-incubated with BubR1_705-1050_ and analyzed by SEC. CENP-E_4E_-GST and BubR1_705-1050_ did not co-elute, indicating they did not bind to each other (Fig 3B). In total, our data indicate the C-terminal helix of BubR1 is necessary but not sufficient to interact with CENP-E_2055-2358_, while the glutamate patch (2313-2319) in CENP-E is essential to support the interaction.

**Figure 3:**
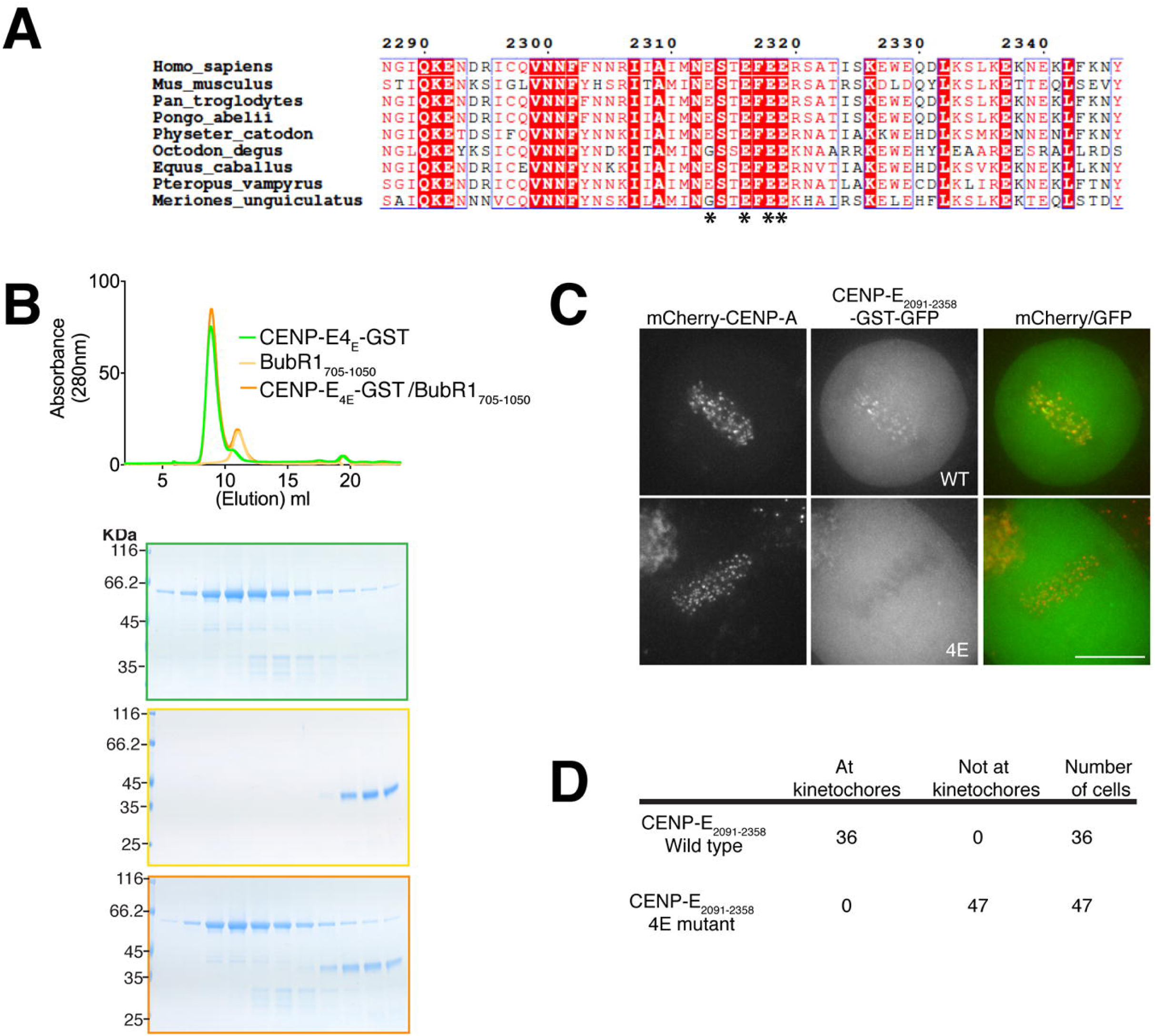
CENP-E uses an acidic patch to bind BubR1. (A) Sequence alignment of the human CENP-E_2287-2246_ with mouse, chimpanzee, orangutan, sperm whale, degu, horse, flying fox and gerbil. Boxed red and blue are the conserved and similar amino acids across all species, respectively. Amino acids in red are those with conserved properties in at least 3 sequences. The glutamates necessary for BubR1 binding in CENP-E are marked with a star (*). (B) Top, SEC analysis and elution profile for CENP-E_4E_-GST (green), BubR1_705-1050_ (yellow) and CENP-E_4E_-GST/BubR1_705-1050_ (orange). Bottom, Coomassie-stained gels showing elution profiles for the corresponding protein complexes. (C) Representative images of live HeLa cells expressing mCherry-CENP-A transfected with CENP-E_2091-2358_-GST-GFP and CENP-E_4E_-GST-GFP constructs. Scale bar: 10 μm. (D) Quantification of the targeting to kinetochores for CENP-E_2091-2358_-GST-GFP and CENP-E_4E_-GST-GFP.

Since potential parallel pathways targeting CENP-E to kinetochores seem to co-exist, we then tested whether CENP-E_2091-2358_-GST-GFP could target to kinetochores in cells and whether this recruitment was only dependent on BubR1. Transiently transfected CENP-E_2091-2358_-GST-GFP is dimeric due to GST and robustly targeted to all kinetochores, however it did not cause chromosome misalignment (Fig 3C), unlike CENP-E_2055-2608_. Thus the minimal kinetochore-targeting domain of CENP-E is unlikely to act as a dominant negative at kinetochores. To test whether CENP-E_2091-2358_-GST targeting to kinetochores is specifically dependent on the glutamate patch mediating interaction with BubR1, we generated CENP_2091-2358_-E_4E_-GST-GFP and hypothesized it should not be able to target to kinetochores. Indeed, CENP CENP_2091-2358_-E_4E_-GST-GFP was not recruited to kinetochores (Fig 3C, D). These data indicate that we have identified the minimal kinetochore-targeting region of CENP-E. Since BubR1 mediates the kinetochore localization of only one pool of CENP-E (see below), this suggests that other interaction partners of CENP-E require the same interaction surface, although we cannot rule out other domains of CENP-E may influence other kinetochore receptors.

The contribution of BubR1 to CENP-E recruitment to kinetochores remains highly controversial (Ciossani et al., 2018; Johnson et al., 2004; Lampson and Kapoor, 2005). Knowing now precisely how CENP-E interacts with BubR1, we next tested to what extent BubR1 contributes to CENP-E localization at kinetochores in mitosis. While CENP-E is highly enriched on unattached, spindle checkpoint-active kinetochores, it is still visible on attached, metaphase kinetochores. To evaluate BubR1’s contribution to CENP-E localization at these distinct kinetochore pools, we used defined synchronization conditions to distinguish the different kinetochore pools. In cells that had been treated with the proteasome inhibitor MG132 for 2.5 hours to enrich for attached, spindle checkpoint-silenced kinetochores, both BubR1 and CENP-E were visible at clear, albeit modest levels in control cells (Fig 4A, S4A). Cells depleted of BubR1 displayed a near complete loss of CENP-E from kinetochores in this situation, suggesting that CENP-E localization to microtubule-attached kinetochores is dependent on the residual pool of BubR1 retained at metaphase chromosomes (Fig 4A-C, Fig S5 A,B). This was also the case when Bub1, essential for the recruitment of BubR1 (Johnson et al., 2004), was depleted (Fig S5A, B). CENP-E localizes to the outer corona of chromosomes, which forms preferentially on unattached kinetochores. To test whether the corona proteins are required for CENP-E kinetochore-targeting on attached kinetochores, too, we depleted the RZZ component ZW10 involved in corona formation (Fig S5). ZW10 depletion, which prevents corona assembly on unattached kinetochores, did not affect CENP-E levels at attached kinetochores under our conditions. These data indicate that while CENP-E can localize to the outer corona, the RZZ complex is not a CENP-E recruiting-factor to kinetochores, as shown previously (Pereira et al., 2018). In the absence of endogenous BubR1, we then expressed full-length BubR1 or BubR1_1-1030_ under an inducible promoter and quantified the corresponding endogenous CENP-E at kinetochores (Fig 4A-I). At aligned kinetochores, CENP-E levels were reduced by half in the presence of BubR1_1-1030._ CENP-E in the presence of BubR1_1-1030_ was however higher than when BubR1 was depleted (Fig 4A-C), suggesting BubR1_1-1030_ enables some low levels of CENP-E to kinetochores. When MG132-arrested cells were treated with a short (5 minutes) pulse of the microtubule-depolymerizing drug nocodazole, a method that has been previously used to test the recruitment of spindle checkpoint components to kinetochores (Vleugel et al., 2015), BubR1 depleted or BubR1_1-1030_ expressing cells were deficient in CENP-E recruitment (Fig 4D-F). After a longer nocodazole treatment (60 minutes), equal levels of CENP-E were observed on BubR1-depleted alone and BubR1_1-1030_-expressing kinetochores (Fig 4G-I), consistent with the idea that BubR1 facilitates initial CENP-E recruitment to SAC-active kinetochores but is not strictly required for CENP-E localization to this subset of kinetochores. In the absence of BubR1, chromosomes are unable to form stable end-on attachments (Fig 4J, K) because of the absence of BubR1-recruited PP2A-B56 (Foley et al., 2011; Kruse et al., 2013; Suijkerbuijk et al., 2012). When BubR1_1-1030_ was expressed in the absence of BubR1, most chromosomes were still able to form a metaphase plate, consistent with the idea that PP2A-B56 targeting was restored in this construct. However, in comparison to cells expressing GFP-BubR1_WT_ we observed a significant increase in cells with misaligned chromosomes (Fig 4K). In these cells, a small number of chromosomes were unable to congress and displayed high levels of GFP-BubR1_1-1030_ at kinetochores, indicating spindle checkpoint activation (Fig 4J, K). This phenotype is very similar to that of CENP-E depletion or knockout suggesting that a pool of CENP-E required for efficient chromosome alignment was missing (Fig 1D) (Schaar et al., 1997). CENP-E, however, was present on the same kinetochores, presumably through a pathway that does not depend on the C terminus of BubR1. The BubR1 C-terminal helix specifically recruits one pool of CENP-E to kinetochores in mitosis and during spindle checkpoint activation. This interaction seems to be required for the productive chromosome alignment and biorientation of chromosomes. In this situation, the CENP-E recruitment to kinetochores through another pathway does not seem to enable full chromosome alignment. Importantly, in our experiments, the GFP-BubR1 construct was expressed with levels similar to endogenous BubR1 (Fig 4L). Overall our data indicate that BubR1 recruits CENP-E specifically to bioriented chromosomes and is important for rapid recruitment of CENP-E to unattached kinetochores during SAC activation. Yet another pathway also promotes CENP-E localization to kinetochores in the absence of BubR1 and Bub1 during the maintenance of SAC.

**Figure 4:**
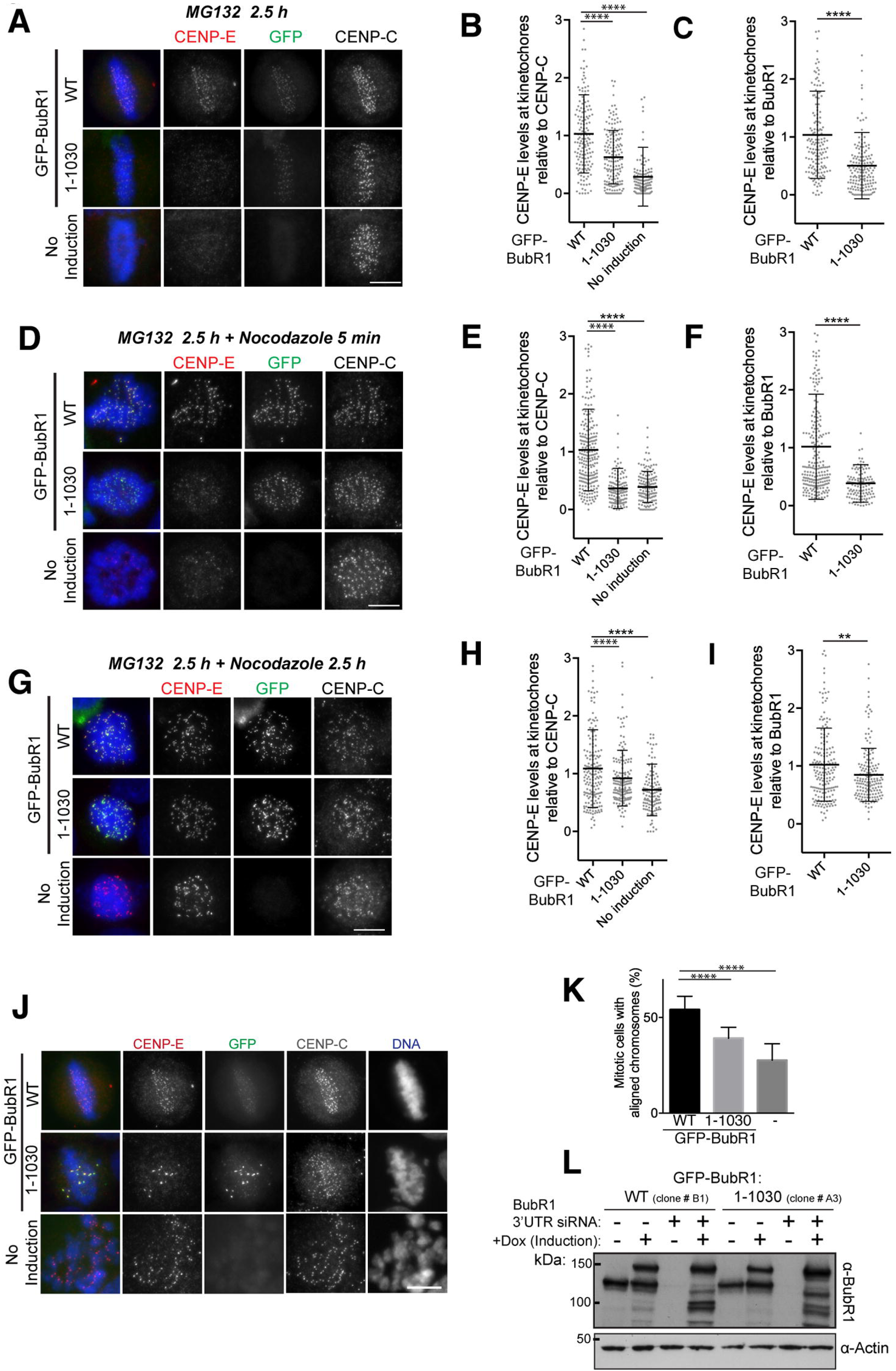
BubR1-dependency of CENP-E recruitment to kinetochores and chromosome alignment. (A) Representative immunofluorescence images of HeLa cells treated with BubR1 siRNA and induced to express GFP-BubR1 WT and GFP-BubR1_1-_ 1030, stained with CENP-E, CENP-C and Hoechst after treatment with MG132 for 2.5 hrs. Scale bar: 10 μm. (B, C) Scatter plots showing CENP-E intensity relative to CENP-C and GFP-BuBR1 at individual kinetochores plotted as grey circles, with mean and standard deviation represented by black lines. Measurements were carried out across 2 independent experiments with cells expressing either GFP-BubR1 (n_kinetochores_= 154) or GFP-BubR1_1-1030_ (n_kinetochores_= 162), and cells with endogenous BubR1 depleted but no GFP induced (n_kinetochores_= 135). For CENP-E:BubR1 ratio, a Student’s T-test was performed, with **** indicating P < 0.0001. (D) Same as in (A). Cells were treated with MG132 for 2.5 hrs and nocodazole for 5 minutes. (E, F) Scatter plots showing CENP-E intensity relative to CENP-C and GFP-BuBR1 at individual kinetochores plotted as grey circles, with mean and standard deviation represented by black lines. Measurements were carried out across 2 independent experiments with cells expressing either GFP-BubR1 (n_kinetochores_= 228), or GFP-BubR1_1-1030_ (n_kinetochores_= 133), and cells with endogenous BubR1 depleted but no GFP induced (n_kinetochores_= 140). For CENP-E:BubR1 ratio, a Student’s T-test was performed, with **** indicating P < 0.0001. (G) Same as in (A). Cells were treated with MG132 and nocodazole for 2.5 hrs. (H, I) Scatter plots showing CENP-E intensity relative to CENP-C and GFP-BubR1 at individual kinetochores plotted as grey circles, with mean and standard deviation represented by black lines. Measurements were carried out across 2 independent experiments with cells expressing either GFP-BubR1 (n_kinetochores_= 176), or GFP-BubR1_1-1030_ (n_kinetochores_= 166), and cells with endogenous BubR1 depleted but no GFP induced (n_kinetochores_= 131). For CENP-E:BubR1 ratio, a Student’s T-test was performed, with **** indicating P < 0.0001. (A, D, G) Conducted across 2 experiments with 5-10 cells measured per condition. (J) Representative immunofluorescence images of HeLa cells treated with BubR1 siRNA and induced to express GFP-BubR1 WT and GFP-BubR1_1-1030_, stained with CENP-E, CENP-C and Hoechst after treatment with MG132 for 2 hrs. (K) Graph showing percentage of cells with at least 1 misaligned chromosome for BubR1-depleted cells induced to express GFP-BubR1, BubR1_1-1030_ or without induction. Error bars represent standard deviation. (L) Western blot for cells in Figure 5 probed for BubR1 and actin as a loading control.

CENP-E is an essential motor, targeting to unattached kinetochores and playing a critical role in the congression, maintenance and biorientation of chromosomes (Shrestha and Draviam, 2013; Vitre et al., 2014; Wood et al., 1997). Here we show that BubR1 is a nanomolar affinity partner of CENP-E in mitosis and we reveal the molecular basis for the CENP-E-BubR1 interaction. The formation of a CENP-E-BubR1 complex is not dependent on post-translational modifications. Similarly to Ciossani et al, our work indicates the pseudokinase domain of BubR1 associates with the C-terminal kinetochore-targeting domain of CENP-E. However they used a construct, which also has the centrosome-targeting domain (Fig S1C) and a second microtubule-binding site (Ciossani et al., 2018; Gudimchuk et al., 2013). In this study, we precisely map the domain of CENP-E necessary for kinetochore targeting, separating it from its centrosome and microtubule-binding functions. This domain is monomeric and associates with a 1:1 stoichiometry with the pseudokinase domain of BubR1, suggesting that full-length CENP-E can associate with 2 molecules of BubR1 at one time with nanomolar affinity. Given the high local concentration of BubR1 at unattached kinetochores, this creates an avidity effect for CENP-E binding to kinetochores to promote its recruitment. Thus the affinity of CENP-E for BubR1 may even be higher than that *in vitro*. There could also be cooperative binding of BubR1 to CENP-E. Our data reveal BubR1 relies on its divergent and basic C-terminal helix for CENP-E binding, creating a unique and specific association to the mitotic motor. Yet this helix is not sufficient. On CENP-E a small acidic patch is critical to specify the interaction with BubR1. Mutation of these amino acids prevents the targeting of this CENP-E_2055-2358_ domain to kinetochores and consequently compromises chromosome alignment.

Previous work on how CENP-E localizes to kinetochores remains unclear and the extent to which it requires BubR1 is conflicting. It is likely due to experimental differences between protocols. Indeed we show that BubR1 facilitates the rapid and initial recruitment of CENP-E to kinetochores at the onset of SAC signaling. Once the SAC is on for a significant period of time, we found CENP-E levels become identical at kinetochores in the presence or absence of BubR1 in good agreement with previous work (Ciossani et al., 2018). Our data indicate that BubR1 is a major interactor of CENP-E at kinetochores but there are distinct yet redundant pathways to recruit CENP-E. BubR1 increases the kinetics of CENP-E recruitment to kinetochores during spindle checkpoint activation. The other pathways contribute to a slower but robust targeting of CENP-E to kinetochores. However they are not sufficient to restore the CENP-E function in chromosome alignment and biorientation. The BubR1-dependent recruitment of CENP-E to kinetochores is therefore essential for correct alignment and biorientation of kinetochores. In the absence of this CENP-E pool at kinetochores, the kinetochore-microtubule attachment is compromised, even when high levels of CENP-E are present (Fig 4J, K). Our work to reveal the molecular basis for BubR1/CENP-E binding at kinetochores will now further facilitate the identification of BubR1-independent pathways that allow CENP-E to associate with kinetochores and the outer corona, to define the contribution of this CENP-E pool to chromosome alignment and biorientation. Future work is needed to address the nature and regulation of the multiple interaction partners of CENP-E to understand its critical role in chromosome congression and maintaining biorientation of kinetochores.

## Author contributions

TL and JW designed the project. TL, DH, JW, EAB, CS and AGK performed experiments TL, DH, EAB, AGK, UG and JW performed data analysis and interpretation. JR secured funding for mass spectrometry equipment. JW wrote the manuscript. All revised the manuscript.

## Acknowledgements

The inducible Cas9 sgCENP-E RNA cell line was a generous gift from Iain Cheeseman. pFL MultiBac His-BubR1:Bub3 was a kind gift from A. Musacchio. J.R. is supported by the Wellcome Trust Senior Research Fellowship (Grant 103139). J. W. is supported by a Wellcome Senior Reseach Fellowship (207430). DH was supported by an MRC Senior Non-Clinical Fellowship (MR/K006703/1) awarded to UG. We thank Martin Wear for help with SEC-MALS. The Wellcome Centre for Cell Biology is supported by core funding from the Wellcome Trust (203149), a Multi-User Equipment grant [101527] for the Edinburgh Protein Production Facility, and Multi-User Equipment grant [108504] for Orbitrap Fusion Lumos.

## Methods

### Cloning

To assay the localization in cell culture of CENP-E subdomains, various constructs were generated from CENP-E transcript variant 1 (NM_001813.2) and cloned into pBABE-puro containing an N- or C-terminal GFP tag and using restriction enzymes (Cheeseman and Desai, 2005). Bacterially-expressed constructs were cloned in pET-3aTr (Tan, 2001). Details of constructs and cloning are listed in Table S2.

### Protein expression, purification and assays

All constructs for bacterial expression were transformed in *E. coli* BL21-CodonPlus (DE3)-RIL. Cultures were induced with 0.5 mM IPTG when OD_600_=0.6 for 4 hours at 25°C or overnight at 18°C for BubR1_705-1050_. Cells were re-suspended in lysis buffer (50 mM HEPES pH 7.5, 500 mM NaCl, 40 mM imidazole, 1 mM EDTA, 5 mM β-Mercaptoethanol) supplemented with 1 mM PMSF and cOmplete EDTA-free protease inhibitor cocktail (Roche) and lysed by sonication. The lysate was cleared by centrifugation (50 minutes, 22,000 RPM) in a JA 25.50 rotor (Beckman Coulter), filtered and loaded onto a HisTrap HP column (GE Healthcare). Proteins were eluted in elution buffer (lysis buffer with 500 mM imidazole). Constructs containing a 3C protease cleavage site were incubated overnight in dialysis buffer (25 mM HEPES pH 7.5, 300 mM NaCl, 10 mM imidazole, 1 mM EDTA, 5 mM β-Mercaptoethanol) with 3C protease and then loaded onto a HisTrap HP column (GE Healthcare). The protein was then concentrated and loaded on a Superdex 200 Increase 10/300 GL (GE Healthcare) pre-equilibrated in size-exclusion chromatography buffer (20 mM HEPES pH 7.5, 300 mM NaCl or 500mM for CENP-E_2055-2608_, 1 mM EDTA, 1mM DTT).

Constructs for insect cell expression were transfected and expressed in SF9 cells using the Bac-to-Bac^®^ expression system (Thermo Fisher Scientific). Expression was carried out for 72 hours at 27°C. Cells were resuspended in lysis buffer supplemented with 1 mM PMSF and cOmplete EDTA-free protease inhibitor cocktail (Roche) and lysed by sonication. The lysate was cleared by centrifugation (60 minutes, 40,000 RPM) in a Type 45 Ti rotor (Beckman Coulter), filtered and loaded onto a HisTrap HP column (GE Healthcare). Proteins were eluted and purified by size-exclusion chromatography as the constructs expressed in bacteria.

Bacteria expressing MBP-BubR1_1031-1050_ were re-suspended in MBP lysis buffer (50 mM HEPES pH 7.5, 500 mM NaCl, 1 mM EDTA, 1 mM DTT) supplemented with 1 mM PMSF and cOmplete EDTA-free protease inhibitor cocktail (Roche) and lysed by sonication. The lysate was cleared by centrifugation (50 minutes, 22,000 RPM) in a JA 25.50 rotor (Beckman Coulter), filtered and loaded onto an MBPTrap HP column (GE Healthcare). Proteins were eluted in elution buffer (lysis buffer with 10 mM Maltose). The fractions containing the protein were concentrated and gel filtered on a Superdex 75 increase 10/300 GL (GE Healthcare) pre-equilibrated in size-exclusion chromatography buffer.

For ITC, CENP-E_2091-2358_-GST was purified in complex with 6His-BubR1_705-1050_. Both lysates were mixed, cleared by centrifugation, filtered and loaded onto a HisTrap HP column. After overnight incubation with 3C protease, the complex was further purified on a Superdex Increase 200 10/300 GL column (GE Healthcare) in separation buffer (50 mM HEPES pH 7.5, 800 mM NaCl, 1 mM EDTA, 1mM DTT). The fractions containing CENP-E_2091-2358_-GST and BubR1_705-1050_ were pooled independently then dialysed against the ITC buffer. All binding assays were carried out on a Superdex Increase 200 10/300 GL column in binding buffer (20 mM HEPES pH 7.5, 150 mM NaCl, 1 mM EDTA, 1mM DTT). Proteins were mixed in equimolar ratio around 7μM.

### Size-exclusion chromatography coupled to multi-angle light scattering (SEC-MALS)

Size-exclusion chromatography (ÄKTA PURE**™**, GE Healthcare) coupled to UV, static light scattering and refractive index detection (Viscotec SEC-MALS 20 and Viscotek RI Detector VE3580, Malvern Instruments) were used to determine the absolute molecular mass of the indicated proteins in solution. 100 µL of CENP-E_2055-2608_ and CENP-E_2055-_ 2358 at 1 mg·ml^-1^ were run on a calibrated Superdex-200 10/300 GL Increase (GE Healthcare) size exclusion column pre-equilibrated in gel filtration buffer (described above) at 22 °C with a flow rate of 1.0 ml·min^-1^. Light scattering, refractive index (RI) and A_280nm_ were analyzed by a homo-polymer model (OmniSEC software, v5.02; Malvern Instruments) using the following parameters: ∂A_280nm_ / ∂c = 0.429 AU·ml·mg^-1^, and 0.530 AU·ml·mg^-1^, CENP-E_2055-2608_ and CENP-E_2055-2358,_ respectively, ∂n / ∂c = 0.185 ml·g^-1^ and a buffer RI value of 1.336.

### Mass spectrometry

Samples were prepared and digested as previously published (McHugh et al., 2019). Following digestion, samples were acidified with 10% TFA until pH<2.5 and spun onto StageTips as described in (Rappsilber et al., 2003). Peptides were eluted in 40 μl of 80% acetonitrile in 0.1% TFA and concentrated down to 1 μl by vacuum centrifugation (Concentrator 5301, Eppendorf, UK). Samples were then prepared for LC-MS/MS analysis by diluting them to 5 μl with 0.1% TFA. LC-MS-analyses were performed on a Q Exactive mass spectrometer (Thermo Fisher Scientific, UK) coupled on-line, to an Ultimate 3000 RSLCnano System (Dionex, Thermo Fisher Scientific, UK). Peptides were separated on a 50 cm EASY-Spray column (Thermo Scientific, UK) assembled in an EASY-Spray source (Thermo Scientific, UK) and operated at 50°C. Mobile phase A consisted of 0.1% formic acid in water while mobile phase B consisted of 80% acetonitrile and 0.1% formic acid. Peptides were loaded onto the column at a flow rate of 0.3 μl min^-1^ and eluted at a flow rate of 0.2 μl min^-1^ according to the following gradient: 2 to 40% buffer B in 90 min, then to 95% in 11 min and 2 to 40% in 120 min and then to 95% in 11 m. FTMS spectra were recorded at 70,000 resolution (scan range 400-1400 m/z) and the ten most intense peaks with charge between 2 and 6 of the MS scan were selected with an isolation window of 2.0 Thomson for MS2 (filling 1.0E6 ions for MS scan, 5.0E4 ions for MS2, maximum fill time 60 ms, dynamic exclusion for 50 s).

The MaxQuant software platform (Cox and Mann, 2008) version 1.5.2.8 was used to process raw files and the search was conducted against *Homo sapiens* complete/reference proteome set of Uniprot database (released in February, 2016), using the Andromeda search engine (Cox et al., 2011). The first search peptide tolerance was set to 20 ppm while the main search peptide tolerance was set to 4.5 pm. Isotope mass tolerance was 2 ppm and maximum charge to 7. Maximum of two missed cleavages were allowed. Carbamidomethylation of cysteine was set as fixed modification. Oxidation of methionine and acetylation of the N-terminal were set as variable modifications.

### Circular Dichroism

Circular dichroism (CD) spectra in the far-ultraviolet region (185-260 nm) for CENP-E_2055-_ 2358 and CENP-E_2055-2608_ (0.1mg/mL) in CD buffer (10 mM Potassium phosphate pH 7.5, 200 mM NaF, 0.5 mM DTT) were recorded using a CD spectrometer (Jasco-J-810) at 10°C (1 mm path length quartz cell). Data was analysed using DichroWeb (http://dichroweb.cryst.bbk.ac.uk, Whitmore and Wallace, 2008).

### Isothermal titration calorimetry (ITC)

Isothermal titration calorimetry (ITC) experiments were carried out to determine the affinity and stoichiometry of the *BubR1*/*CENP-E* complex. BubR1_705-1050_ and CENP-E_2091-_ 2358-GST were extensively dialysed into ITC buffer (20 mM HEPES pH7.5, 150 mM NaCl, 1 mM EDTA, 0.005% Tween-20, 0.5 mM TCEP); prior to the experiment to minimise heats of dilution upon titration. Protein concentrations were determined by absorption at 280 nm; extinction coefficients ε for BubR1_705-1050_ and CENP-E_2091-2358_-GST were 63370 M^-1^cm^-1^ and 62800 M^-1^cm^-1^ respectively. 276 μM BubR1_705-1050_ was titrated into 205.4 µl of 25 μM CENP-E_2091-2358_-GST at 37 °C in 11 aliquots: 1 of 0.5 μl followed by 10 x 3.8 μl. The reference power was set to 3 μcal/s and syringe rotation 750 rpm. The enthalpy of binding was analysed with correction for heat of dilution using the software package provided by the instrument manufacturer (Auto-iTC200 microcalorimeter; Malvern Instruments). Data were fit to a simple binding model with one set of sites.

### Low-angle rotary shadowing and electron microscopy

CENP-E_2055-2608_ at a concentration of 100 µg/mL in gel filtration buffer with 30% glycerol were sprayed onto a mica sheet (TAAB). CENP-E_2055-2608_ was shadowed with 2.5 nm of platinum at 5° angle and 9 nm of carbon using Leica EM ACE600. Replicas were detached in water and placed on non-coated grids (Type 400 mesh, TAAB). Images were obtained using a JEM-1400Plus transmission electron microscope (JEOL) operated at 90 kV. Electron micrographs were acquired using GATAN OneView camera.

### Cell Culture and experiments

HeLa cells were used and maintained in DMEM (Lonza) supplemented with 5% CO_2_ at 37°C in a humidified atmosphere. The inducible HeLa Cas9 sgRNA CENP-E cell line was obtained from Iain Cheeseman (McKinley and Cheeseman, 2017) and maintained in a tet-free medium. Cells are monthly checked for mycoplasma contamination (MycoAlert detection kit, Lonza). Transient transfections were conducted using Effectene reagent (Qiagen) according to the manufacturer’s guidelines. GFP-BubR1 Wild Type (WT) and 1-1030 HeLa cell lines (#B1 and #A3 respectively) were generated with single integrated copies of the desired transgenes using the T-Rex doxycycline-inducible Flp-In system, and were chosen for equal expression levels as seen by immunofluorescence and western blotting.

GFP-BubR1 was induced 6 h before a 48-h siRNA depletion of endogenous BubR1 using oligonucleotides against the 3′ UTR (5′-GCAATCAAGTCTCACAGAT-3′, (Espert et al., 2014). A second induction was performed 24 h into the siRNA depletion. 26.5 h prior to fixing cells were subjected to Thymidine arrest for 16 h followed by a 10.5 h release. For the final 2.5 h, MG132 was added at 20 µM to increase the metaphase population.

### CRISPR Cas9 knockout

To induce Cas9 expression, cells were treated with 1 µg/ml doxycycline (Sigma Aldrich) for 48-72h, changing the medium with fresh doxycycline every 24h to induce the knockout.

### Microscopy

For live-cell imaging, HeLa cells were imaged in Leibovitz L15 media or DMEM (Life Technologies) supplemented with 10% FBS + penicillin/streptomycin (Gibco) at 37°C using a Deltavision core microscope (Applied Precision) equipped with a CoolSnap HQ2 CCD camera. 4-10 z-sections were acquired at 0.5 µm steps using a 60x objective lens. For immunofluorescence, cells were washed with PBS and fixed by one of two methods, either fixed in cold methanol for 10 minute at −20°C and then permeabilised with cold acetone for 1 minute at –20°C, or pre-extracted with 0.4% Triton-X in PHEM for 1 min and then fixed with 3.8% formaldehyde in PHEM buffer (60 mM Pipes, 25 mM HEPES, 10 mM EGTA, 2 mM MgSO_4_, pH 7.0) for 20 minutes. For experiments with HeLa Flipin inducible cells, fixation was performed with PTEMF (20 mM Pipes-KOH, pH 6.8, 0.2% Triton X-100, 1 mM MgCl_2_, 10 mM EGTA, and 4% formaldehyde) for 12 min. Immunofluorescence in human cells was conducted using antibodies against mouse □-tubulin (Sigma, 1:1000), mouse CENP-E (Abcam, Ab5093, 1:1000 or 1:200), rabbit Centrin (kind gift from I. Cheeseman, 1: 1000), guinea pig CENP-C (pAb, MBL PD030, 1:2000) and human ACA antibodies (Cambridge Biosciences, 1:100). Hoechst 33342 (ThermoFisher Scientific; H3570) was used to stain DNA. For experiments with small molecule inhibitors, ZM447439, MLN8237 and nocodazole were used at a final concentration of 2 □M, 300 nM and 300 ng/ml, respectively for 2h. A widefield Eclipse Ti2 (Nikon) microscope equipped with a Prime 95B Scientific CMOS camera (Photometrics) was used for imaging. Z-sections were acquired at 0.2-µm step size. Images were stored and vizualised using an OMERO.insight client (OME) (Allan et al., 2012). Mean kinetochore fluorescence intensity within a circular ROI with a 10 pixel diameter was measured for with a background intensity recorded in an adjacent cytoplasmic area. Relative CENP-E and BubR1 value for each kinetochore are calculated by subtracting the background values and dividing them by the background corrected ACA/CENP-C signal for that kinetochore. Data was analyzed using ImageJ (Schneider et al., 2012).

### Statistics and reproducibility

Statistical analyses were performed using GraphPad Prism 6.0. No statistical method was used to predetermine sample size. All experiments were performed and quantified from at least three independent experiments, unless specified and the representative data are shown.

### Data availability

All data and reagents supporting the findings of this study are available from the corresponding author on request.

## Figure legends

**Supplementary figure 1:** (A) Scatter dot plot showing the quantification of CENP-E intensity normalized to ACA (logarithmic scale) in prometaphase cells, metaphase cells and cells treated with nocodazole. Each point represents the intensity of CENP-E over ACA at one kinetochore. Black line represents the median. (B) Representative immunofluorescence images of HeLa cells for quantification in (A). (C) Representative immunofluorescence images of HeLa cells transfected with GFP-CENP-E_2260-2608_ and stained for kinetochores (ACA), centrioles (centrin2) and DNA (Hoechst). Scale bar: 10 μM. (D, E) Scatter dot plots (logarithmic scale) showing quantification of GFP-CENP-E_2055-2608_ fluorescence intensity at kinetochores in the presence and absence of endogenous CENP-E (knockdown using inducible CRISPR-Cas9) for non polar kinetochores (D) and polar kinetochores (E). Each point represents the intensity of GFP-CENP-E_2055-2608_ at one kinetochore. Black line represents the median. Asterisks indicate ordinary unpaired T-test significance value. ****P<0.0001

**Supplementary figure 2:** (A) Elution profile (black line, left y-axis) from a size-exclusion chromatography (SEC) run with subsequent multi angle light scattering (MALS) analysis for CENP-E_2055-2608_ (top) and CENP-E_2055-2358_ (bottom). Outcome of the MALS analysis for the peak is presented in blue (molecular weight, right y-axis). (B) Table showing the predicted and measured mass, stoichiometry of the proteins and polydispersity index (Mw/Mn) values. (C) Circular dichroism spectra for 100 μg/ml CENP-E_2055-2358_ (orange) and CENP-E_2055-2608_ (black) indicating the proteins are predominantly α-helical. (D) Table summarizing the secondary structure features determined from the circular dichroism spectra in C. (E) Representative CENP-E_2055-2608_ particles observed after rotary shadowing. The double arrow shows the length of one particle (40 nm).

**Supplementary figure 3:** CENP-E kinetochore-targeting domain associates with the pseudokinase domain of BubR1. (A) Schematic diagram showing the identification of CENP-E_2055-2358_ interacting proteins. (B) Mass spectrometry table of proteins identified to specifically associate with CENP-E_2055-2358_, reporting the number of peptides identified and molecular weight of protein partners. (C, D) Top, SEC analysis and elution profile for the indicated constructs for CENP-E (green) and BubR1 (yellow) and CENP-E/BubR1 (orange). Bottom, Coomassie-stained gels showing elution profiles for the corresponding protein complexes.

**Supplementary figure 4:** (A) Top, SEC analysis and elution profile for CENP-E_2055-2358_ (green), BubR1_705-1050_ (yellow) and CENP-E_2055-2358_/BubR1_705-1050_ (orange). Bottom, Coomassie-stained gels showing elution profiles for the corresponding protein complexes. (B) Top, SEC analysis and elution profile for GST (green), BubR1_705-1050_ (yellow) and GST/BubR1_705-1050_ (orange). Bottom, Coomassie-stained gels showing elution profiles for the corresponding protein complexes, showing GST does not interact with BubR1.

**Supplementary figure 5:** (A, C) Representative immunofluorescence images of HeLa cells treated with indicated siRNA and treated with MG132 for 2.5 hours with and without 5 minutes nocodazole treatment. Cells were stained with CENP-E, BubR1, CENP-C and Hoechst. Scale bar: 10 μm. (B, D) Scatter dot plot showing the quantification of CENP-E intensity normalized to CENP-C. Numbers of kinetochores analyzed for cells treated with 2.5h MG132 and 2.5h MG132+ 5 minutes nocodazole after RNAi depletion were respectively: n _control_= 185, 140; n_BuBR1_=140, 98; n_Bub1_=140, 145; n_ZW10_=111, 159; n_Bub1/BubR1_=120, 100; n_Bub1/BubR1/ZW10_=60, 100. Asterisks indicate significance value performed using a ANOVA one-way test. **** indicate a P-value<0.0001.

**Supplementary Table S1**: list of proteins found after CENP-E_2055-2358_ pulldown by mass spectrometry

**Supplementary Table S2**: list of constructs and cloning details used in this paper

