## Supplementary figures and images for "The C-terminal helix of BubR1 is essential for CENP-E-dependent chromosome alignment"

### Supplementary Figure 1

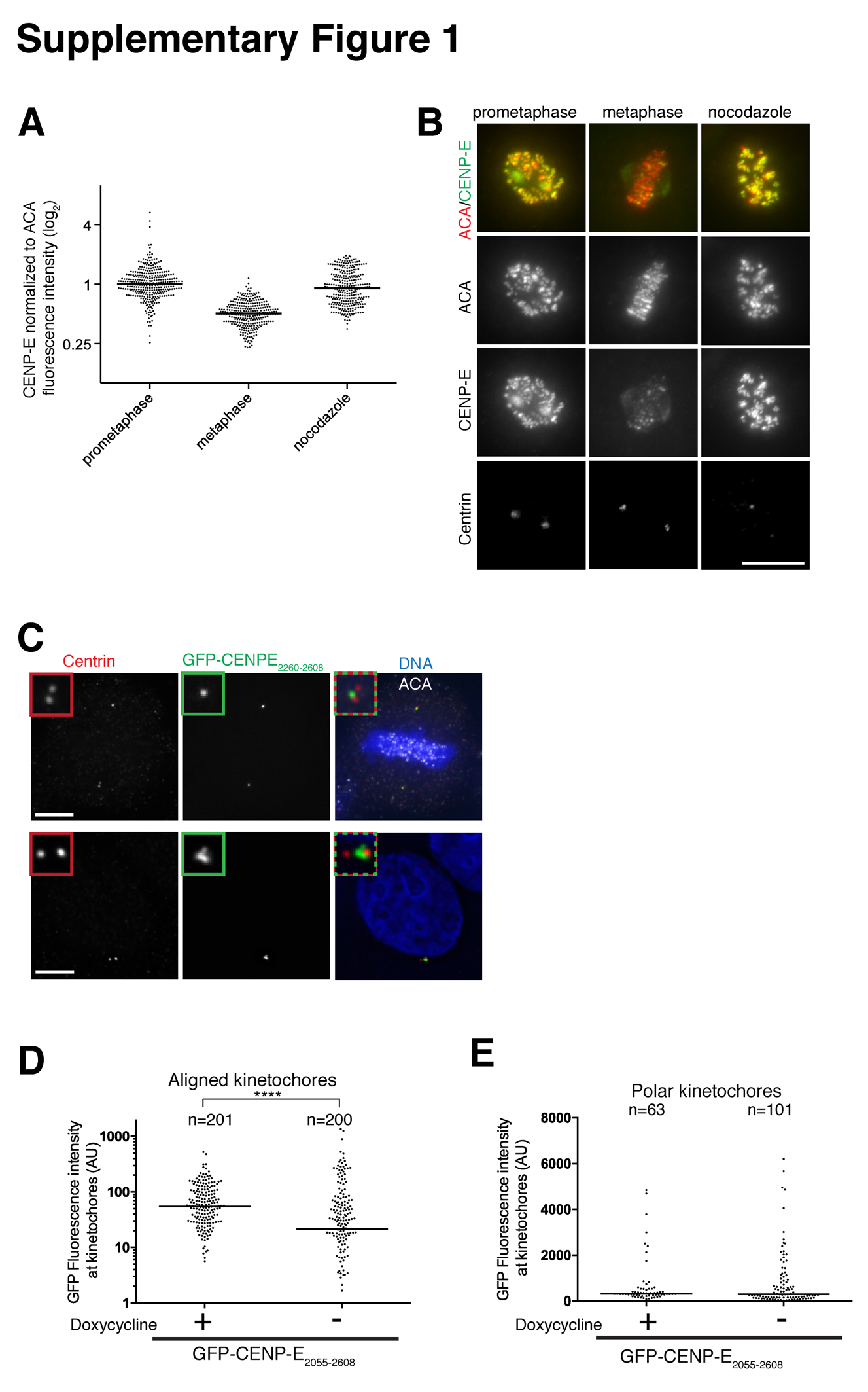

### Supplementary Figure 2

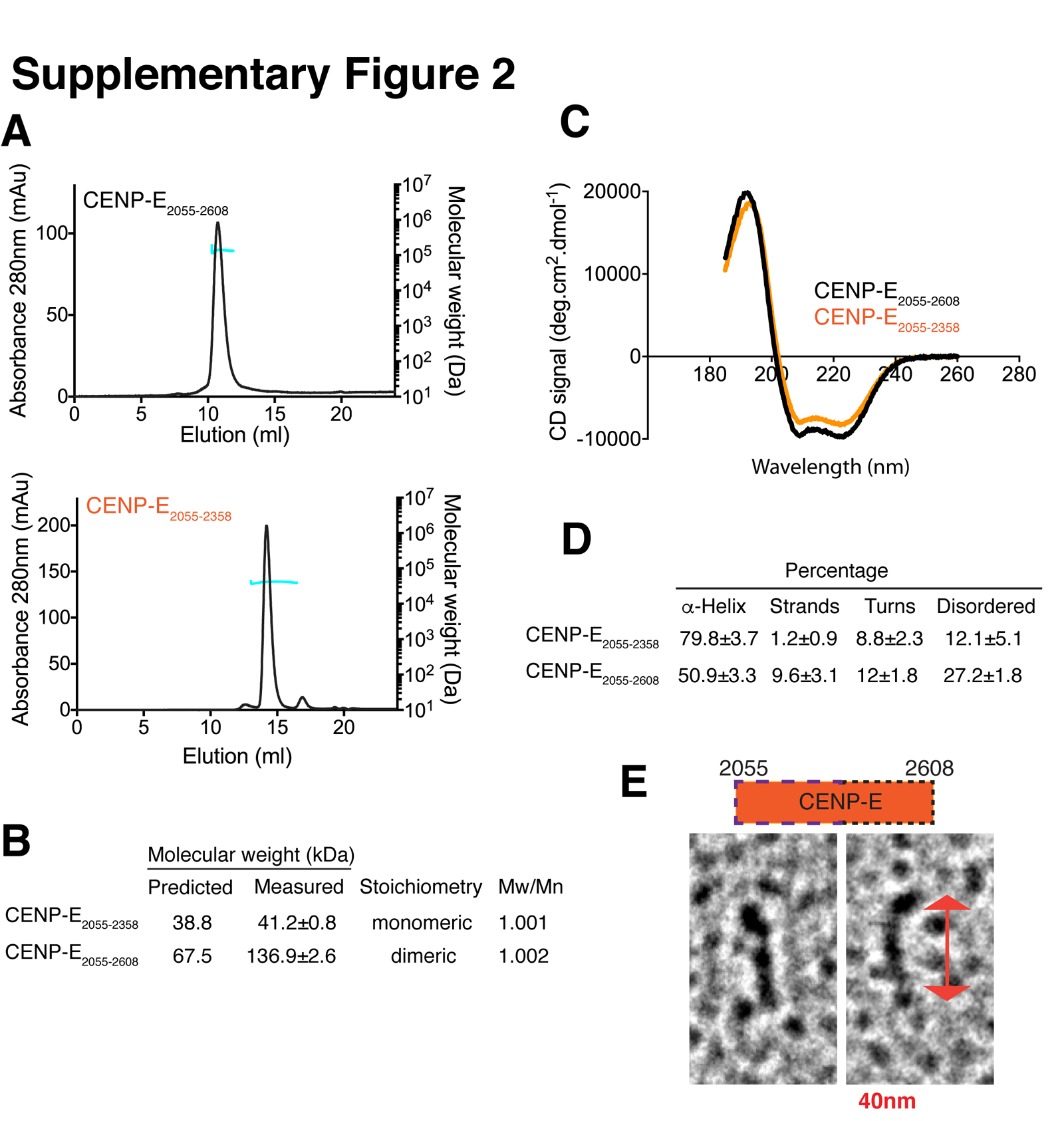

### Supplementary Figure 3

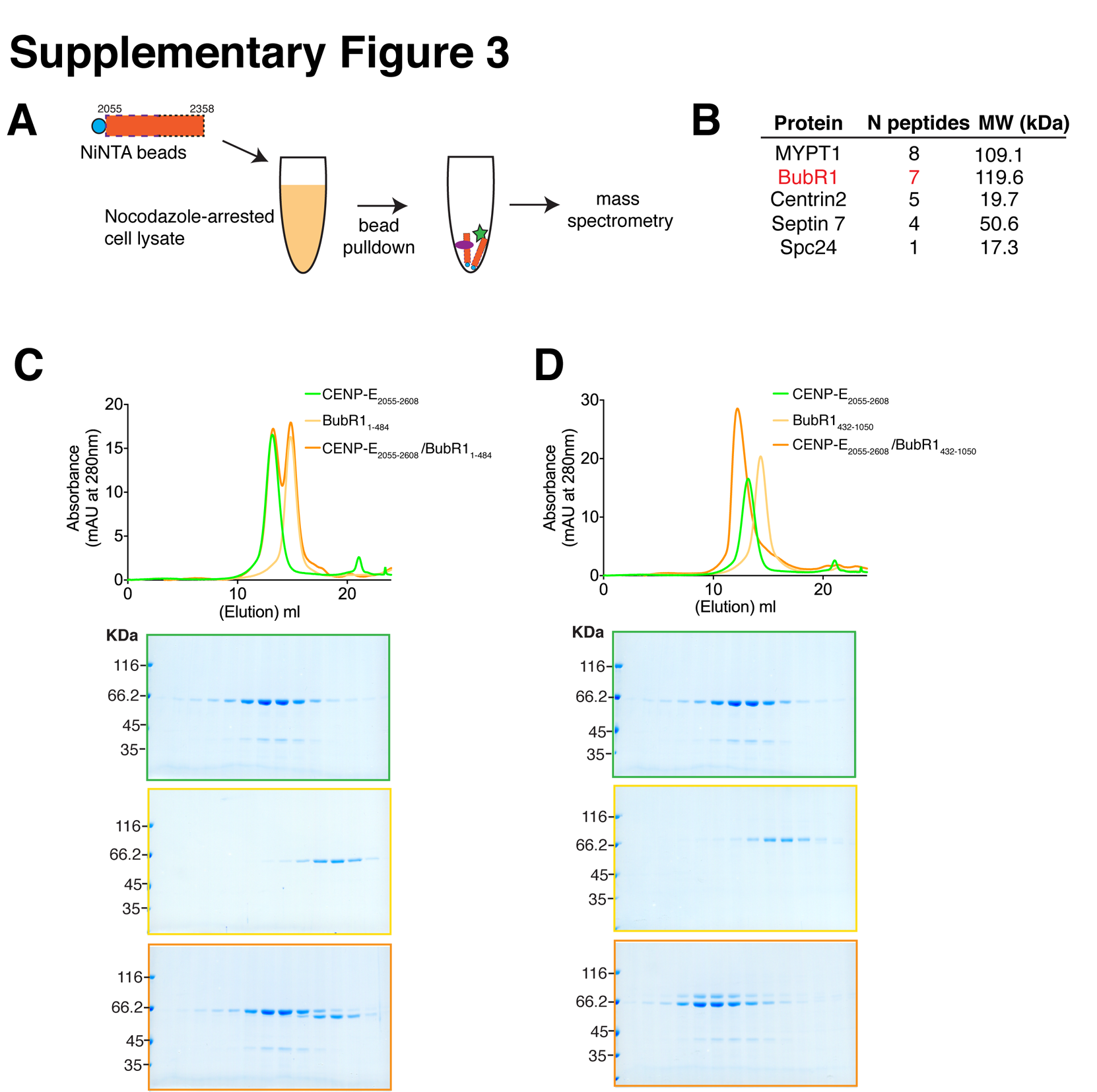

### Supplementary Figure 4

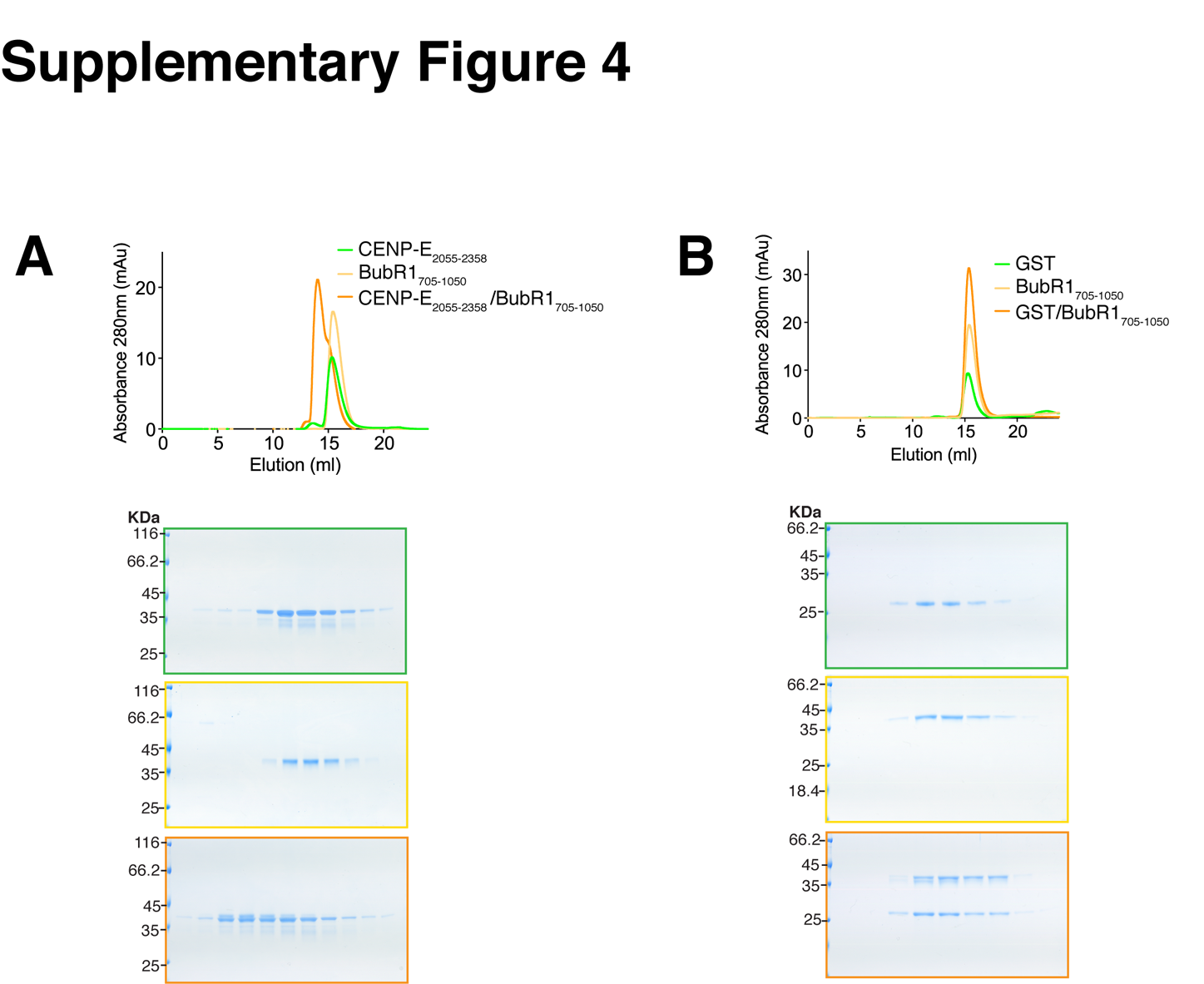

### Supplementary Figure 5

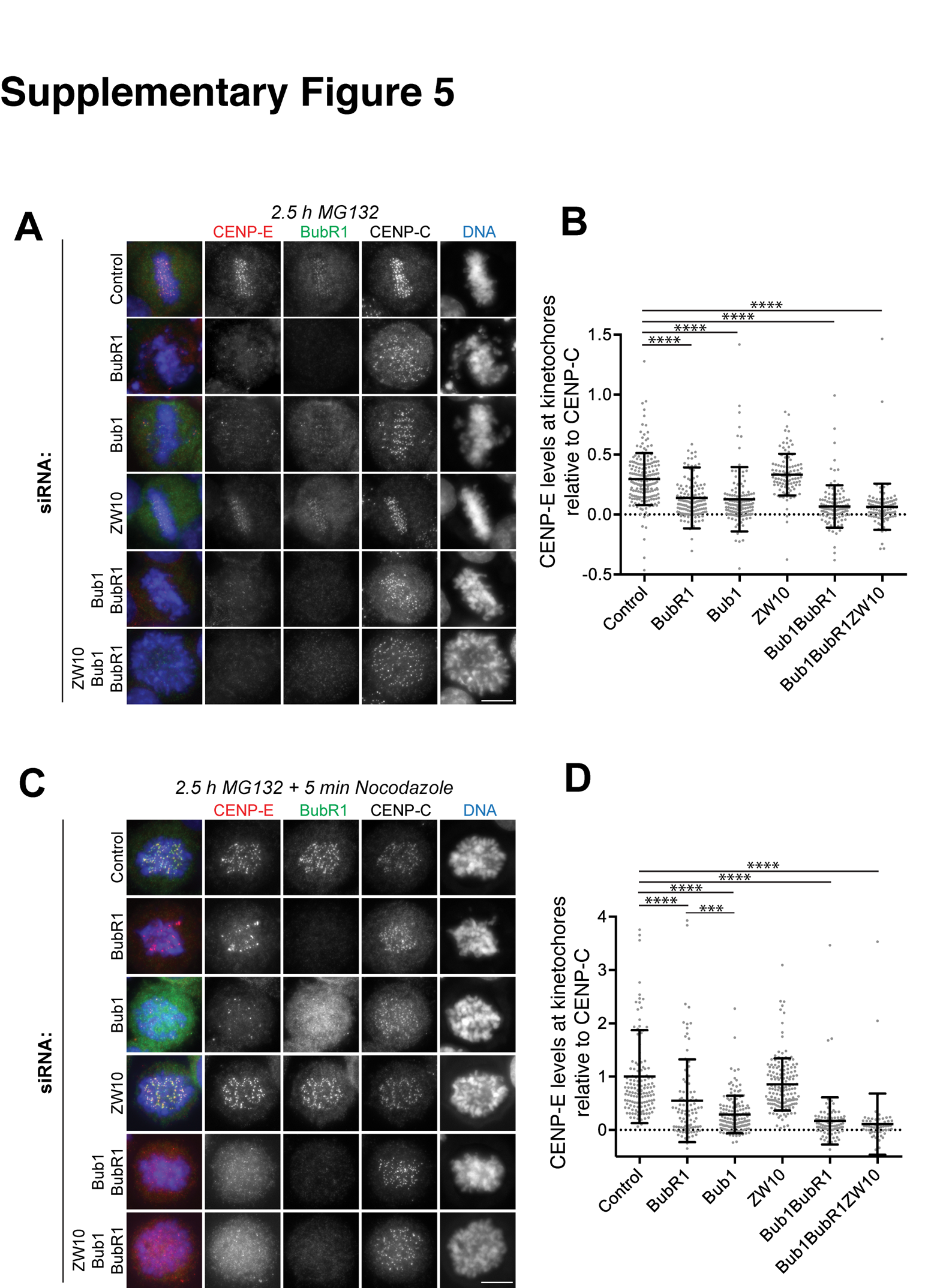
